# Chromosome 17: neXt-MP50 C-HPP Challenge Completed

**DOI:** 10.1101/734657

**Authors:** Hongjiu Zhang, Omer Siddiqui, Yuanfang Guan, Gilbert S. Omenn

**Affiliations:** Department of Computational Medicine and Bioinformatics, University of Michigan, Ann Arbor, Michigan 48109, United States; Department of Electrical Engineering and Computer Science, University of Michigan, Ann Arbor, Michigan 48109, United States; Department of Internal Medicine, University of Michigan, Ann Arbor, Michigan 48109, United States; Department of Human Genetics and School of Public Health, University of Michigan, Ann Arbor, Michigan 48109, United States; Microsoft, 555 110th Avenue NE, Bellevue, Washington 98004, United States

**Keywords:** neXt-MP50, protein existence evidence levels (PE), mass spectrometry, protein-protein interactions

## Abstract

The C-HPP Team for Chromosome 17 has documented meeting the neXt-MP50 Missing Protein Challenge. Protein-level PE1 evidence including MS and protein-protein interaction data from the global proteomics community has reduced the number of PE2,3,4 proteins coded on Chromosome 17 by 50 since the 2016 baseline in neXtProt. As a follow-up to our analysis and prioritization of Chr 17 missing proteins last year, we describe what predictions and guidance were successful in this update and we explain the dynamics of a net reduction of PE2,3,4 Missing Proteins. This analysis can serve as a guide for other C-HPP chromosome teams seeking high-quality evidence for the 2129 remaining missing proteins.

## INTRODUCTION

Chromosome 17 had 148 PE2,3,4 Missing Proteins (MPs) in October 2016 when C-HPP announced the neXt-MP50 Challenge for each chromosome team to advance 50 MPs to PE1 status, with neXtProt release 2016-01 as the baseline. In 2018, we performed an extensive analysis on the progress from the global proteomics community that identified 43 Chr 17 missing proteins, showing the contributions of mass spectrometry (MS) (25), protein-protein interactions (PPI) (12), MS + PPI (3), and disease mutation, biological characterization, or PTMs (3) to the net reduction of Chr 17 MPs to 105 in neXtProt 2018-01^1^. From the progress of the past year, as documented in the neXtProt release 2019-01, Chr 17 has 98 remaining MPs, thus achieving the neXt-MP50 goal with highly confident identification of a net of 50 PE2,3,4 Missing Proteins per chromosome since 2016. Of course, attention must turn to the remaining MPs. As was true when presented to the C-HPP Workshop in Saint-Malo, France, in May 2019, this analysis is intended as a model for detailed assessment of the progress for each of the chromosomes, including others that have now achieved the MP50 target^2^.

## RESULTS FOR CHR 17 DURING 2018

Here we report the complex path to a net of 7 fewer PE2,3,4 MPs for Chr 17 during 2018 for neXtProt 2019-01. Figure 1 shows the dynamics underlying this net reduction of 7 Chr 17 MPs during 2018-2019 to reach the total of 50 fewer Chr 17 MPs since 2016.

**Figure 1.**
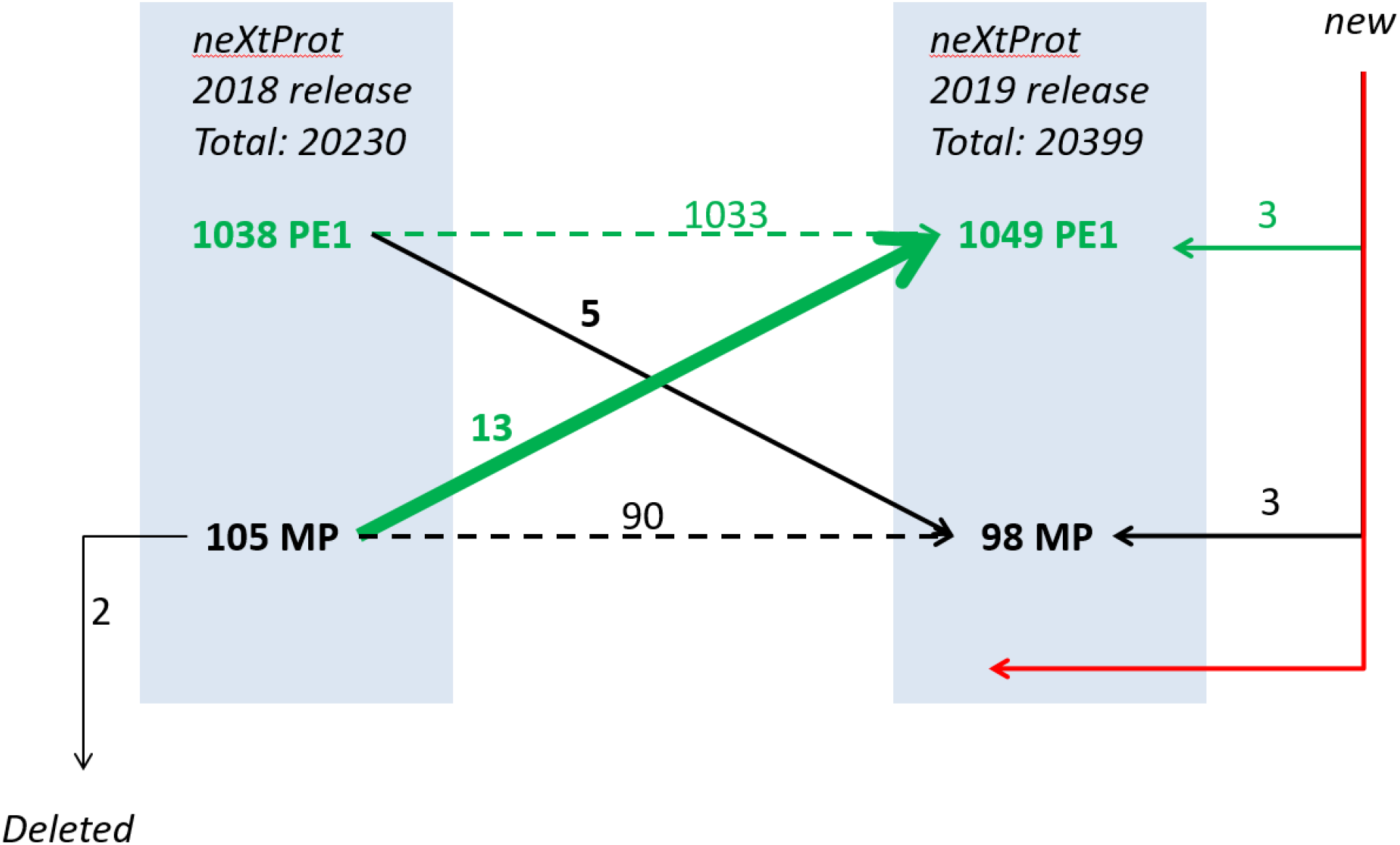
The dynamic changes in neXtProt for Chr 17 during the year from 2018-01 to 2019-01, showing promotion of PE2,3,4 missing proteins to PE1 high-confidence expressed proteins, as well as new entries to neXtProt and demotion and deletion of previous PE1 and PE2,3,4 proteins.

Table 1A shows the 11 PE2, the 1 PE3 (KRTAP9-7), and the 1 PE4 (RNF222) MPs promoted to PE1 in 2018. For all decisions based on MS, full documentation of the uniquely mapping, non-nested peptides of 9 or more amino acids in length is present in PeptideAtlas 2019-01. We list the number of such peptides and the specimen(s) analyzed. In some cases, one or more peptides are from different specimens (see PeptideAtlas).

**Table 1.** Changes in the PE1 protein class and the PE2,3,4 Missing Proteins class between 2018-01 and 2019-01 versions of neXtProt database

| *A. 13 Chr 17 Missing Proteins promoted to PE1 from PE2,3,4 during 2018* |  |  |  |
| --- | --- | --- | --- |
| **neXtProt ID** | **Gene name** | **Type of Evidence**<br>**(# Non-Nested Unique**<br>**Peptides in**<br>**PeptideAtlas >= 9 aa)** | **Major Tissue Source** |
| NX_P29275 | ADORA2B | MS (2) | HeLa, Placenta |
| NX_Q8IVW1 | ARL17B | MS (2) | Melanoma |
| NX_A6NF36 | CCDC182 | MS (5) | Testis |
| NX_Q86WR6 | C17orf64 | MS (3) | Testis |
| NX_A8MZ36 | EVPLL | MS (2) | Saliva, esophagus |
| NX_Q9UQ05 | KCNH4 | MS (2) | HeLa |
| NX_Q9BYQ8 | KRTAP4-9 | MS (2) | Plasma |
| NX_A8MTY7 | KRTAP9-7 | MS (2) | Plasma |
| NX_Q9ULX5 | RNF112 | IHC | Glioblastoma cells |
| NX_A6NCQ9 | RNF222 | MS (2) | Embryonic and induced pluripotent stem cells |
| NX_O15375 | SLC16A5 | MS (4) | Colon, HeLa |
| NX_Q6DHY5 | TBC1D3G | PPI | Y2H with ZRANB1 |
| NX_Q6UXU6 | TMEM92 | MS (2) | HeLa |

| **B. 3 new Chr 17 PE1 entries from outside** |  |  |  |
| --- | --- | --- | --- |
| **neXtProt ID** | **Gene name** | **Type of Evidence**<br>**(#Unique Peptides in PeptideAtlas)** | **Major Tissue Source** |
| NX_A0A0K2S4Q6 | CD300H | Immunoprecipitation | Myeloid lineage cells |
| NX_A0A1B0GVT2 | SMIM36 | MS (2) | Retina |
| NX_A0A1B0GUW6 | SPEM3 | MS (>10) | Sperm |

| **C. 5 Chr 17 Proteins demoted from PE1 to PE2,3,4 MPs from 2018-01 to 2019-01** |  |  |
| --- | --- | --- |
| **neXtProt ID** | **Gene name** | **# Unique Peptides**<br>**in PeptideAtlas** |
| NX_Q8N9I5 | FADS6 | 0 |
| NX_O14610 | GNGT2 | 1 |
| NX_Q07627 | KRTAP1-1 | 0 |
| NX_Q9BYP8 | KRTAP17-1 | 1 |
| NX_O14894 | TM4SF5 | 1 |

| **D. 3 New Chr 17 PE2,3,4 MPs from outside** |  |
| --- | --- |
| **neXtProt ID** | **Gene name** |
| NX_Q53H64 | ANKRD40CL (PE4) |
| NX_A6NIN4 | NF227 (PE2) |
| NX_A0A1B0GWK0 | PVALEF (PE3) |
| ***E. 2 Chr 17 MPs removed*** |  |
| **neXtProt ID** | **Gene name** |
| NX_Q9Y5M1 | FAM215A (PE2) |
| NX_O15453 | NBR2 (PE2) |

Among these 13 MPs newly promoted to PE1, 11 are based on MS, 1 on PPI, and 1 on immunohistochemistry (IHC). EVPLL, SLC16A5, TMEM92, and RNF222 are proteins we highlighted with potential stranded peptide spectra^1^ (Table 1), *i.e.*, they already had a singleton qualifying peptide in PeptideAtlas and neXtProt and a second we found in ProteomeXchange; they are now classified as PE1 based on additional spectral evidence in PeptideAtlas. Among the 11 promoted from PE2 to PE1, we also had identified C17orf64 and RNF112 as priority candidates because there was already one proteotypic peptide in PeptideAtlas; they both, plus ADORA2B, had high transcript TPM values^1^. TMEM92 and KRTAP9-7 were prioritized based on PPI, but then identified by MS, suggesting that, though we prioritized KRTAPs and membrane proteins for PPI based on difficulty in solubilizing membrane proteins and high sequence similarities for KRTAPs, it is still fruitful to look for them in MS data.

Some trends stand out in the MS data for these 13 promoted MPs. CCDC182 and C17orf64 both had proteotypic peptides found in the testis. These MS experiments used a multiple protease strategy; many of the peptides found came from LysC and GluC digestion in addition to trypsin^3^. EVPLL and RNF222 each had a peptide found in the same experiment on saliva, which employed a combination of LysC and trypsin^4^. The use of multiple proteases serves to produce more proteotypic peptides that can be detected by MS, which is especially important for proteins that have relatively few sites for trypsin digestion. Another salient point is the use of nontraditional samples for detecting remaining missing proteins, such as saliva, esophagus (the second EVPLL peptide), colon (SLC16A5), and cancers (melanoma for two peptides for ARL17B). Interestingly, 5 different experiments with Hela cells are represented in Table 1A, suggesting HeLa cells as an area of potential for finding more MPs, a trend which was not noted when previously analyzing the 2018 release of neXtProt for Chromosome 17. However, a large-scale reanalysis of 41 publicly-available HeLa cell proteomics datasets found only one missing protein (NX_O75474, chr 10); altogether there are 116 HeLa datasets in PRIDE^5^.

Meanwhile, Table 1B shows 3 new neXtProt entries assigned to PE1: CD300H, SMIM36, and SPEM3. CD300H was detected by immunoprecipitation; it is not found in PeptideAtlas, because it lacks MS data. SMIM36 has two uniquely-mapping peptides and is classified as canonical in PeptideAtlas, citing Pinto et al^6^; it is not clear why it was not curated as PE1 in UniProt/Swissprot/neXtProt until 2018. SPEM3 is canonical in PeptideAtlas with multiple published detections in sperm and testis and a total of 39 distinct peptides.

In Table 1C are 4 PE1 proteins demoted by neXtProt to PE2 plus 1 (KRTAP17-1) demoted to PE3. These apparently reflect re-curation in UniProt/Swissprot adopted by neXtProt. In PeptideAtlas, each has either zero (FADS6 and KRTAP1-1) or 1 (GNGT2, TM4SF5, KRTAAP17-1) unique peptide. FADS6 is not found in PA, while KRTAP1-1 has 22 observed peptides, but not 2 uniquely-mapping peptides, so the entry is labeled “representative”. Those with 1 peptide are classified as weak or marginally distinguished. Apparently, these were all previously PE1 based on non-MS data that were reassessed.

In Table 1D, former PE2 entries NBR2 (next to BRCA 1 gene) and FAM215A (uncharacterized apoptosis-related protein) were removed, apparently, reflecting reassessments of the quality of the non-protein evidence. Neither has a current entry in neXtProt and neither has any observed peptides in PeptideAtlas, despite predicted observable peptides.

Finally, in Table 1E, there are 3 new MPs for Chromosome 17: RNF227 (PE2) and ANKRD40CL (PE4) were promoted from PE5, while PVALEF is a brand-new PE3 entry; none has a unique peptide from MS. RNF227 is a putative ring finger metal ion binding protein also described in neXtProt as long intergenic non-protein coding RNA 2581, observed only as a transcript. ANKRD40CL is an ANKRD40 C-terminal-like 114 aa protein with a single 8 aa peptide; its entry is based on prediction. PVALEF is parvalbumin-like EF-hand-containing protein regulating muscle contraction, with no proteomics evidence and no entry in PeptideAtlas, based on homology.

For the past year, that makes a net of 7 fewer PE2,3,4 MPs for Chr 17, as shown in Figure 1. So Chr 17 has 98 PE2,3,4 MPs and 1049 PE1 proteins in neXtProt release 2019-01.

neXtProt is certainly a dynamic database, in part due to proteogenomics being a dynamic field, with reannotations of the genome and the proteome in the interconnected databases.

## CONCLUSION

Chromosome 17 has achieved the neXt-MP50 Challenge goal of confidently detecting 50 PE2,3,4 Missing Proteins from the reports of the global community. Chromosomes 19, 1, and 5 have also reached this target as of neXtProt 2019-01^2^. Similar analyses investigating the use of MS/PPI and different tissue samples by or for the other chromosome teams could accelerate recognition of progress across the entire human proteome^2^.

## ACKNOWLEDGMENTS

This work is supported by NIH grants P30ES017885 and U24CA210967 (G.S.O.) and NSF 1452656 (Y.G.).

